# On the Validity of the Saccharum Complex and the Saccharinae Subtribe: A Re-assesment

**DOI:** 10.1101/2020.07.29.226753

**Authors:** Dyfed Lloyd Evans, Shailesh Vinay Joshi

## Abstract

The ‘Saccharum Complex’ represents an hypothetical collective of species that were supposedly responsible, through interbreeding, for the origins of sugarcane. Though recent phylogenetic studies have cast doubt on the veracity of this hypothesis, it has cast a long shadow over the taxonomics of the Andropogoneae and the Saccharinae subtribe. Though evidence suggests that Saccharum s.s. is comprised of only three true species, according to Kew’s GrassBase there are as many as 34 species in *Saccharum s.l*. Our recent work has shown that many of these species are millions of years divergent from Saccharum. As the Saccharum complex represents the species that sugarcane breeders attempt to introgress into sugarcane, and as the Saccharinae, in its current form, covers almst 12 million years of Andropogoneae evolution an update on the extents of the Taxonomic and customary groupings is much needed. Based on the latest sequence based phylogenies and the inclusion of traditional taxonomics we develop an integrated view of the Saccharinae + Saccharum complex species in the context of the major groupings within the Andropogoneae. We use this phylogeny to re-circumscribe the limits of both the Saccharinae subtribe and the Saccharum complex group of interbreeding species.

## Introduction

The Saccharinae subtribe is part of the Andropogoneae tribe of the PACMAD clade of Poaceae (true grasses). The Andropogoneae were defined by Dumortier (1824). Though the Saccharinae was first coined in 1815 by Knuth (1815). However, his definition was deemed invalid and, formally the Saccharinae were defined by Grisebach (1846). They type species for the Saccharinae is *Saccharum officinarum* L.

Though the ‘Saccharum complex’ is an informal definition initially intended to represent those species thought to have contributed to the origins of sugarcane, it has had such an undue influence on taxonomics that the origins and development of the concept needs to be understood. Muckergee (1957) first coined the term ‘Saccharum complex’ where he pointed out that the four genera *Saccharum, Erianthus, Sclerostachya* and *Narenga* constituted a closely related inter-breeding group concerned in the origin of sugarcane. Daniels et al. (1975) included *Miscanthus* section *Diandra* to the ‘Saccharum complex’ as it was thought to be involved in the origin of Saccharum. This concept was further refined by Clayton and Renvoize (1986) who extended the subtribe Saccharinae to include the genera: *Erianthus* Michaux, *Eriochrysis* P. Beauv., *Eulalia* Knuth, *Eulaliopsis* Honda, *Homozeugos* Stapf, *Imperata* Crillo, *Lophopogon* Hack, *Microstegium* Nees, *Miscanthus* Andersson, *Pogonatherum* P. Beauv., *Polliniopsis* Hayata, *Polytrias* Hack, *Saccharum* and *Spodiopogon* Trin. (as a result many of these genera have been re-classified as *Saccharum* and genus *Saccharum* now comprises between 35 and 40 species, mostly from the tropics and sub-tropics). The suggestion being, that all these genera are closely allied to *Saccharum* and were actually involved in the evolution of sugarcane’s ancestors. This paper has had considerable taxonomic influence and, for example, both the New World and Old World genera of *Erianthus* as well as *Narenga porphyrocoma* are now all included within *Saccharum sensu lato*.

This definition has also significantly influenced the delimeting of genus *Saccharum* itself, with many authorities also treating *Saccharum* in a broader sense (*Saccharum sensu lato*). For example, Kew’s GrassBase currently recognizes 36 species within *Saccharum* (http://www.kew.org/data/grasses-db/sppindex.htm#S) and Tropicos presents 189 distinct species names under the *Saccharum* genus (https://www.tropicos.org/name/Search?name=Saccharum) though many of these names are synonyms. Indeed, the circumscription of *Saccharum* remains highly controversial and has changed significantly over the past century. Several phenetic studies have indicated strong molecular differentiation between *Saccharum* and *Erianthus* (Besse et al., 1998; Nair et al., 2005; Selvi et al., 2006). Conversely, a phylogenetic analysis based on the internal transcribed spacer (ITS) of the nuclear ribosomal DNA (Hodkinson et al., 2002) found no support for this division, even though this study suggested that *Saccharum s.l.* is polyphyletic. Even the taxonomic delimitation between *Saccharum* and *Miscanthus* is not clear, with intergeneric hybrids occurring between them (Clayton & Renvoize, 1986; Hodkinson et al., 2002).

As the Saccharum complex/Saccharinae comprises the gene pool that sugarcane breeders use when attempting to introgress useful characteristics into sugarcane the true relationship of these genera and species to each other, as determined by molecular techniques is of considerable import and relevance. This is especially the case, as modern molecular techniques do not support the concept of a ‘saccharum complex’ (D’Hont et al. 2008). Moreover, there is increasing evidence that *Saccharum* is a well-defined lineage that diverged over a long evolutionary period from the lineages leading to the New World *Erianthus* and Old World *Miscanthus* genera (Grivet et al. 2004; Estep et al. 2014; Lloyd Evans & Joshi, 2016).

Kellogg (2013) also added *Euclasta, Spathia, Lophopogon* and *Leptatherum* to the Saccharinae. However, the most recent treatment of the Saccharinae is that of Soreng et al. (2017) where the Saccharinae subtribe is circumscribed to include the following genera: *Agenium, Asthenochloa* (introduced as a member of the Sorghinae), *Cleistachne, Erianthus, Eriochrysis* (syn *Leptosaccharum*), *Euclasta* (syn *Indochloa*), *Eulalia, Hemisorghum, Homozeugos, Imperata, Lasiorhachis, Leptatherum* (syn *Polliniopsis*), *Miscanthidium, Miscanthus, Narenga, Polytrias, Pseudodichanthium, Pseudopogonatherum, Pseudosorghum, Saccharum, Sclerostachya, Sorghastrum, Sorghum, Trachypogon, Tripidium, Veldkampia*.

Some of these genera are clearly not closely related to Saccharum. Indeed, recent phylogenetic studies indicate that *Tripidium* is over 11 million years divergent, from *Saccharum* with *Eriochrysis* being even more divergent (Lloyd Evans et al., 2019)). Low copy number phylogenetics indicates that genus *Sorghum* is not monophyletic (Estep et al. 2014, Lloyd Evans et al. 2019) and ITS phylogenetics demonstrates that *Microstegium* is not monophyletic (Snyman et al. 2018). Chloroplast-based phylogenetics diverges from low copy number phylogenetics and ITS-based phylogenetics (Lloyd Evans et al. 2019, Snyman et al 2018) demonstrating that reticulate (network) evolution is commonplace in the Andropogoneae and the Saccharinae. As a result, the current circumscription of the Saccharinae is in dire need of review and updating based on the latest phylogenetics.

Other examples of clearly misplaced taxa are *Saccharum* (*Lasiorhachis*) *hildebrandtii*, which whole chloroplast analysis clearly places within *Sorghum* (Piot et al. 2018). Extended ITS phylogenetics places both *Saccharum hildebrandtii* and *Saccharum perierri* (Lloyd Evans and Hughes 2020) within *Sorghum*.

Other species are not well studied, but are probably the most closely related to Saccharum. These include Narenga, Saccharum longisetosum (syn Erianthus rockii), Miscanthus nepalensis, Miscanthus nudipes, Erianthus fulvus and Narenga fallax. Some workers (Welker et al. 2015) place the South American Erianthus species, as exemplified by the type, Erianthus giganteus (Walter) P. Beauv. within Saccharum, though the case is not yet proven.

It is clear that the extent of the Saccharm complex requires a new circumscription. We present an ITS-based phylogeny that places the species most closely related to Saccharum in their taxonomic context. We also employ a text searching and community based approach to analyze the taxonomic placement of those genera currently placed in Saccharum and develop a consensus phylogeny based on a combination of ITS and nuclear low copy number gene phylogenetics leading to the most comprehensive molecular view of the relationships between purported members of the Saccharinae subtribe developed to date.

## Results

### ITS-based Phylogeny

The ITS-based phylogeny (Figure 1) places the species most closely related to *Saccharum* within their taxonomic context. *Miscanthus* (along with *Pseudosorghum*) forms an outgroup to *Saccharum* and its allies. The remaining species are sister to *Saccharum s.s.* and can be divided into four distinct groupings. The outgrooup for this clade is a novel clade formed from *Narenga, Miscanthus* and *Erianthus* species. Sister to this grouping are the *Erianthus* species from the Americas. Sister to *Erianthus* are the African *Miscanthidium* species and a novel grouping of *Narenga porphyrocoma, Erianthus rockii* and *Miscanthus fuscus*.

**Figure 1:**
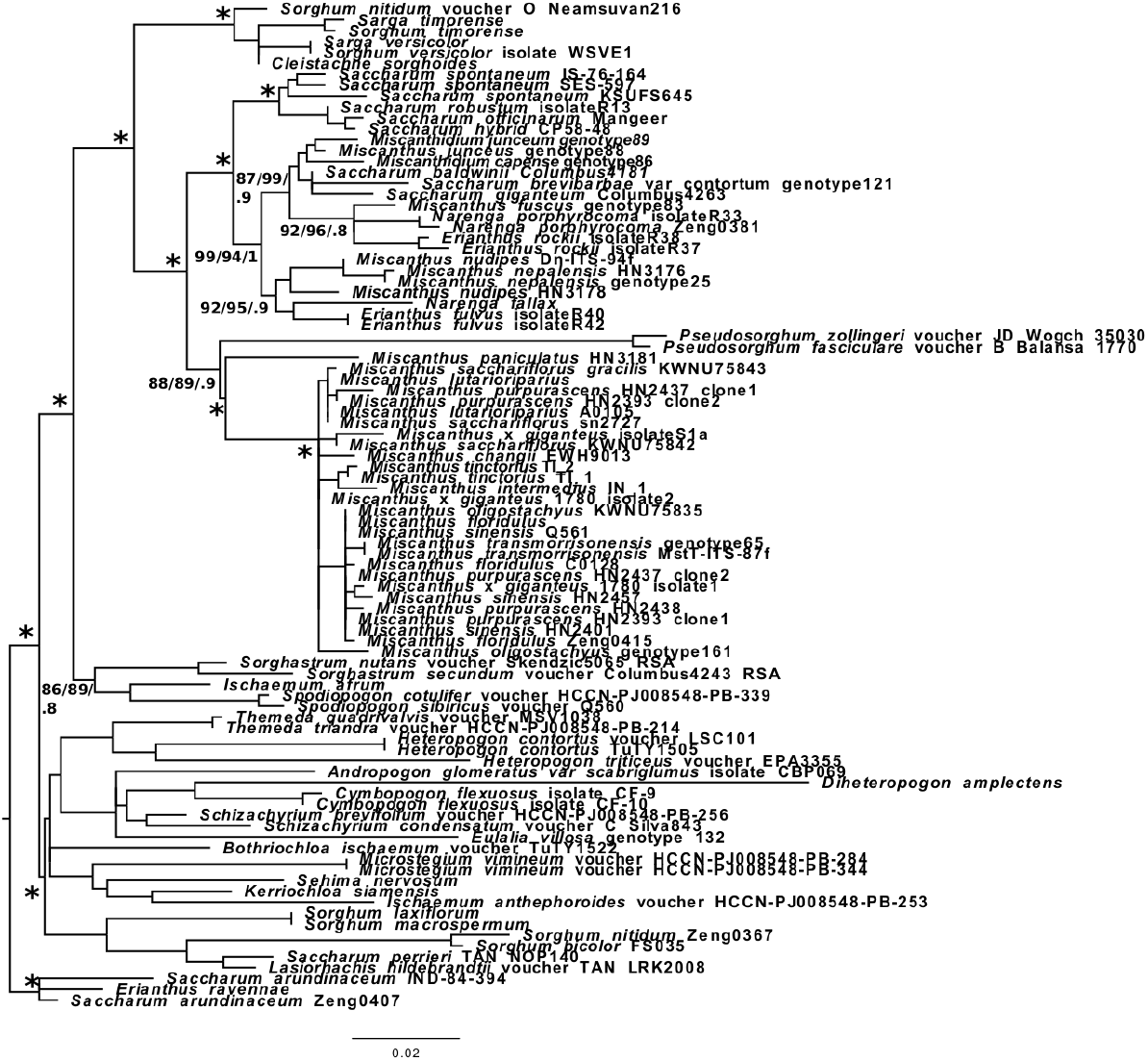
ITS phylogeny of those species and genera most closely related to Sacharum. A phylogeny based on an 600bp ITS (internal transcribed spacer) region encompassing all the species most closely related to Saccharum. This reveals a clade comprised of species with n=15 that is sister to Saccharum. This clade is comprised of Narenga, Miscanthidium, Erianthus (from the Americas) and a novel grouping of Trans-Himalayan species. Support (SH-aLRT, non parametric bootstrap, Bayesian Inference) are given at the main nodes only (* = 100% branch support). The scale bar represents the expected number of substitutions per site.

### Combined Text search, Taxonomic and Phylogenetic Analysis

Each genus purported to lie within the Saccharinae subtribe were taken in tern and the results of natural language processing to derive taxonomic placement and phylogenetic analyses were integrated. The result is the community phy logeny (Figure 2). Using this phylogeny it’s possible to analyze each of the genera purported to lie within the Saccharinae in turn, as below:

**Figure 2:**
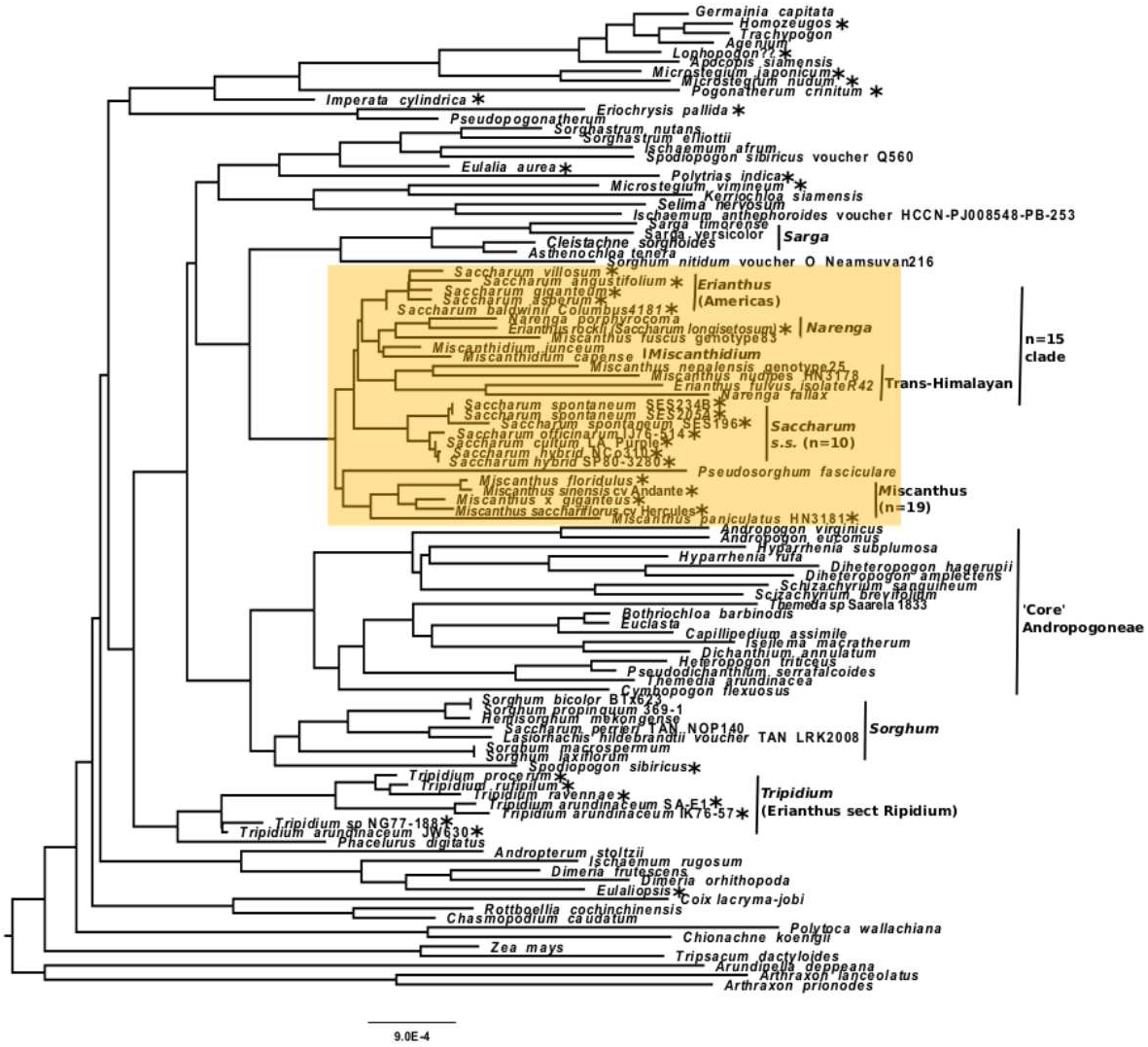
Community phylogeny for the Sacharinae and members of the ‘Saccharum complex’. An integrated phylogeny based on community data from low copy number gene and ITS phylogenies, showing the phylogenetic relationships between all members of the ‘Saccharum complex’. Lineages that could only be placed by traditional Taxonomic means are shown with a dashed line. The shaded box delimits the extent of the core saccharinae and the 3.4 million year window in which wild (non human mediated) hybridization is possible. *Mi crostegium* and *Erianthus* are clearly not monophyletic within this phylogeny. An asterisk (*) represent purported members of the Saccharinae subtribe.

#### Agenium

Guala (1998, 2000), based on ITS phylogenies, places *Homozeugos* as sister to *Tmchypogon* with *Agenium* sister to *Homozeugos* + *Tmchypogon.*

#### Asthenochloa

The placement of *Asthenochloa* has not been widely studied. The most comprehensive analysis (Setshogo 1997) places *Asthenochloa* as sister to *Cleistachne* (which agrees with their cytogenetics as tetraploids with a base chromosome number of 9 and with the pedicellate spikelet absent).

#### Euclasta

*Euclasta* was introduced into the Saccharinae by Soreng et al. (2017) as a member of the Sorghinae. However, the The ITS phylogeny of Skendzic et al. (2007) places *Euclasta* as sister to *Bothriochloa*. This is a member of the Andropogoninae and last shared a common ancestor with *Saccharum*

#### Pseudodichanthium

No taxonomy could be found, but the clossest ally is *Dichanthium* and some workers consider *Dichanthium serrafalcoides* (T.Cooke & Stapf) Blatt. & McCann a synonym for *Pseudodichanthium serrafalcoides* (T. Cooke & Stapf) Bor. *Pseudodicanthium* was removed from *Dicanthium* as *Pseudodichanthium* differed from other *Dichanthium* species in appearance, texture and disposition of the glumes, in the pedicellate spikelet being larger than the sessile, and in the winged glumes.It resembles the genus *Dichanthium* only in the imbricate spikelets of which the lower two or three are homogamous. Tiwar and Chroghe (2019) in their lecotypification of *Pseudodichanthium* placed the genus as closer to *Heteropogon*. However, *Heteropogon* and *Dichanthium* are allies, some 10.3 million years divergent from Saccharum.

#### Pseudopogonatherum

In a whole chloroplast phylogeny, Arthan et al. (2017) placed *Pseudopogonatherum contortum* (as *Eulalia contorta*) within a clade containing *Andropogon burmanicus* and *Parahyparrhenia siamensis* that was sister to *Eriochrysis* and therefore last shared a common ancestor with *Saccharum* about 12 million years ago.

#### Pseudosorghum

Estep et al. (2014) placed *Pseudosorghum* within the *Miscanthus* clade in their low copy number gene phylogeny. However, Arthan et al. (2017) in their whole chloroplast phylogeny placed *Pseudosorghum* as sister to *Eulalia* which would make it 10.1 million years divergent from *Saccharum*. The ITS phylogeny presented in this paper also supports *Pseudosorghum* as being sister to *Miscanthus*. It would therefore appear, as is the case for many Andropogoneae arose as the result of reticulate evolution, with differing genomic and plastome signals it may be appropriate to retain it within the Saccharinae.

#### Veldkampia

The taxonomic position of *Valdekampia* is uncertain, though the original authors tentatively placed this single-species genus within *Saccharum* (Ibaragi & Kobayashi 2007). The reduction of the pedicellate spikelet to just the pedicel hints that this genus might be related to *Cleistachne*, though karyotype information would be required for confirmation.

#### Spathia

Morphological studies place *Spathia* in a clade with *Eulalia* and *Germainia* (Kellogg and Watson 1992). However, recent molecular evidence does not support monophyly of and *Eulalia*+*Germainia* clade. If *Spathia* is closer to *Germainia*, then it is not within the Saccharinae. If it is more closely allied to *Eulalia* then it belongs to a group that is sister to the core Saccharinae.

#### Erianthus

Typically, it is members of *Erianthus* sect *Ripidium* that are included in the Saccharum complex and not *Erianthus* as a whole. However, recently sufficient data has become available to examine the three main branches of *Erianthus*: *Erianthus* sect *Ripidium, Erianthus* species from the Americas (currently part of *Saccharum sensu lato*) and the trans-Himalayan *Erianthus* species (as exemplified by *Erianthus rockii* [syn *Saccharum longisetosum* and *Erianthus fulvus*]). Our recent analysis of *Erianthus* sect *Ripidium* demonstrated that *Erianthus* is polyphyletic and that members of *Erianthus* sect *Ripidium* more properly belong to genus *Tripidium* which is 12 million years divergent from *Saccharum* and is more closely related to *Phacelurus* (a genus of African and Eurasian grasses), (Lloyd Evans et al. 2018). *Erianthus* species from the Americas form a distinct clade that is sister to African *Miscanthidium* species and *Narenga* (including *Erianthus rockii*. The trans-Himalayan species (including *Erianthus fulfus* form an outgrop to all the other clades (Figure 1, Figure 2).

#### Eriochrysis

this genus is even more divergent from *Saccharum* than *Tripidium* and should be excluded from the Saccharum complex as it cannot naturally inter-breed with *Saccharum* (Lloyd Evans et al. 2019). The genus should also be excluded from the Saccharinae.

#### Eulalia

is sister to *Sorghastrum* and forms an outgroup to the core Saccharinae. At over 7 million years divergent from *Saccharum* it must be excluded from the Saccharum complex (Welker et al 2014). However, being in a grouping sister to the Saccharinae it might still be included in this sub-tribe.

#### Microstegium

*Microstegium* is not monophyletic. One part of the genus is related to *Germainia*, whilst *Microstegium vimineum* groups with *Polytrias* and *Eulalia*. More research is needed, but ITS phylogenies place these as the immediate ancestors of the core Saccharinae. However, they are not within the natural hybridization window with *Saccharum* and should be excluded from the core Sacchrinae and the Saccharum complex (Snyman et al. 2018; Lloyd Evans and Hughes 2020).

#### Polytrias

Like *Microstegium*, to which they are related, *Polytrias* species emerge as sister to *Microstegium vimineum* and the core Saccharinae, at least based on chloroplast analyses. However, ITS phylogenies place this genus as more distal to *Saccharum* (Lloyd Evans et al. 2019, Lloyd Evans and Hughes 2020).

#### Imperata

*Imperata* is an ancient hybrid. Its chloroplast phylogeny places it as sister to *Pogonatherum*, thus forming an outgroup to *Sorghum*. However, its genomic sequences (low copy number genes and ITS region) is mch more divergent (at least 14 million years divergent from *Saccharum*) (Estep et al 2014; Lloyd Evans et al. 2018; Lloyd Evans and Hughes 2020).

#### Pogonatherum

Choriplast phylogeny places *Pogonatherum* as sister to *Imperatata* (Lloyd Evans et al. 2019).

#### Polliniopsis

(Now ncluded in *Microstegium*, but see *Microstegium*, above). As no sequence data exists in NCBI for this genus, its position in Figure 1 is captured taxonomically as part of *Microstegium s.s.* and sister to *Apocopis* and *Germainia*.

#### Spodiopogon

Low copy number gene and ITS phylogenies place *Spodiopogon* as ancestral to *Sorghum* and the core Andropogoneae (Estep et al. 2014; Lloyd Evans and Hughes 2020).

#### Eulaliopsis

From whole chloroplast phylogenetic analysis *Eulaliopsis* is sister to *Dimeria* with this grouping last sharing a common ancestor with *Saccharum* 11.6 million years ago (Lloyd Evans et al. 2016) as a result it should be excluded from the Saccharinae and the Saccharum complex.

#### Homozeugos

No sequence data are currently available for this genus of African species. However, work by Gula II (1998) clearly placed *Homozeugos* as sister to *Trachypogon* and did not place this genus within the Saccharinae. *Trachypogon* is sister to *Germainia* and therefore a member of the Germainiinae and not the Saccharinae; the same holds true for *Homozeugos* (Kellogg and Birchler 1993). As ITS regions (NCBI: DQ005006) were available for *Trachypogon plumosus* these were added to the phylogeny of Snyman et al. 2018. The relative position of *Trachypogon* within the phylogeny was mapped to Figure 2 and the sister relationship of *Trachypogon* and *Homozeugos* was captured.

#### Lophopogon

No sequence data are currently available for this genus however, this genus of Indian plants is now placed within the Germainiinae (at least 11 million years divergent from Saccharum).

#### *Sorghum* and *Sarga*

Based on whole chloroplast and chloroplast region phylogenetics, these genera were clustered as *Eusorghum* and *Parasorghum*, respectively. However, low copy number gene analyses and ITS analyses place *Sorghum* as sister to the core Andropogoneae and *Sarga* as sister to the core Saccharinae. When the core Andropogoneae and Saccharinae diverged some 7.5 million years ago hybridization events ocurred between ancestral *Sarga* and *Sorghum* species. *Sarga* gained the *Sorghum* chloroplast type but retained its saccharinae-type genome (Estep et al 2014; Snyman et al 2017, Lloyd Evans et al. 2019, Lloyd Evans and Hughes 2020).

#### Miscanthus

though this genus only diverged from *Saccharum* about 3.4 million years ago (Lloyd Evans and Joshi 2016) is clearly separate and divergent from *Saccharum* (with a base chromosome number of 19 as opposed to 10). It does, however, lie just within the window where wild hybridization with *Saccharum* is possible (Lloyd Evans and Joshi 2016). Chromosome analysis, however, indicate that only certain popyploid forms of *Miscanthus floridulus* are compatible with *Saccharum* hybridization (ref).

#### Miscanthidium

Originally included within *Miscanthus*, there is now broad agreement that *Miscanthidium* forms a distinct and separate genus of African species (Hodkinson 2002). This genus is much more closely related to *Saccharum* (about 2 million years divergent) than *Miscanthus*. Miscanthidium should be included in a new definition of the Saccharinae.

#### Narenga

*Narenga porphyrocoma* is the species that, from chloroplast analyses, is most closely related to Saccharum. Low copy number gene evidence indicates that it hybridized with an ancestral Saccharum species about 2 million years ago (Lloyd Evans et al. 2019). ITS phylogenetics (Figure 1) places this genus within a clade of genera with a base chromosome number of 15. Thus it is a member of the Saccharum complex but should probably be excluded from *Saccharum s.s*.

*Sorghastrum, Kerriochloa, Sehima, Ischaemum, Dimerium* (along with with *Microstegium vimineum* and *Polytrias indica* all form a clade (in low copy number gene phylogenies) that is sister to the Saccharinae. Whether these are members of the Saccharinae (or forma a separate subtribe separate from it) is a matter of debate. What is clear is that these genera are more closely related to *Saccharum* than the majority of the genera described above, however these genera were never included in the Saccharum complex.

## Discussion

Based on the phylogenies presented in this paper we can place all the purported members of the Saccharum complex in their proper phylogenetic position. Given a 3.4–4.2 million year window where hybridization between members of the Andropogoneae is possible in the wild (Lloyd Evans and Joshi 2016) (shaded in Figure 2), this means that the genera *Tripidium* (*Erianthus* sect *Ripidium*), *Eriochrysis, Imperata, Polliniopsis, Homozeugos, Lophopogon* and *Microstegium* (but see below for *Microstegium vimineum*) can be excluded from the Saccharum complex as they cannot naturally inter-breed with *Saccharum. Eulalia* can be excluded as it is sister to the core Andropogoneae and it is generally held that the core Andropogoneae form the division between species that are part of the Saccharinae and those which are not (Kellogg 2013). The positons of *Microstegium vimineum, Pogonatherum* and *Imperata* are more unclear. Both chloroplast and nuclear phylogenetics place *Microstegium viminium* as an outgroup to the core Saccharinae (but outside the wild hybridization window) whilst the exact phylogenetic positions of *Pogonatherum* and *Imperata* as complex ancient hybrids requires more work. What can be said definitely is that they should be excluded from the Saccharum complex, but their position as ancestral to the Saccharinae remains in question. Indeed, Comparing low copy number gene phylogenies with whole chloroplast and ITS phylogenies reveals that may of these genera are complex reticulate hybrids and that further analysis is required to elucidate the true taxonomic placement of these genera. However, taking nuclear phylogenies as representing the ‘true’ phylogenetic placement, what is clear is that they are neither members of *Saccharum* nor members of the Saccharum complex.

The revelaton that Sarga species are an hybrid and that their nuclear phylogenetic signals place them as a natural outgroup to the Saccharinae is a major finding (Estep et al. 2014, Snyman et al. 2018, Lloyd Evans and Hughes 2020). Thus, a genus that was not even considere as being closely related to Saccharum emerges as being more closely related than ¾ of the puroprted members of the Saccharum complex.

Taxonomically, *Sarga* species (and this includes *Cleistachne sorghoides* are the most distal member of the Saccharinae subtribe. Within the core Saccharinae we have *Miscanthus, Miscanthidium, Narenga, Erianthus*, a trans-Himalayan grouping and *Saccharum* itself (Figure 1), (Lloyd Evans and Hughes 2020).

*Pseudosorghum* emerges as sister to *Miscanthus* (Lloyd Evans and Hughes 2020) and should also be included in the Saccharinae.

Thus, the Saccharinae subtribe has a true biological meaning and can be confirmed to contain at least four core genera, with the exact positioning of the two clades within Erianthus requiring further work (though they are part of the Saccharinae). As an outgroup, *Sarga, Asthenochloa* and *Cleistachne* should also be included within the Saccharinae whilst the potential inclusion of *Microstegium vimineum, Pogonatherum* and *Imperata* will require further study.

Our ITS-based phylogeny (Figure 1) positions several new species within the Saccharinae (see Figure 2 for a legend). We have a novel clade that is sister to *Saccharum sensu stricto*. This clade includes *Erianthus, Narenga, Miscanthidium* and a novel clade that contains Trans-Himalayan species and which warrants further investigation. Interestingly, the base chromosomal count for the majority of this group is 15 (Jensen et al. 1989; Sreenivasan & Sreenivasan 1989; Hoshino & Davidse 1988). The exception being the trans-Himalayan outgroup with a base chromosome count of 10 (Mehra & Sharna 1975). There is some evidence that many of the members of this clade are themselves hybrids (Lloyd Evans et al. 2018; Lloyd Evans & Hughes 2020). Thus the different base chromosome number and separate hybrid origins of this grouping would seem to exclude them from *Saccharum*. As such, *Saccharum sensu stricto* includes only those species within genus *Saccharum* itself. Though these species should all be included within the Saccharinae subtribe.

From Figure 2, as well as *Saccharum sensu stricto* and the n=15 clade, the Saccharinae should also include *Miscanthus* and *Sarga* (which also includes *Cleistachne* and *Asthenochloa*).

There is a clade that is sister to the core Saccharinae formed from *Sorghastrum, Ischaemum, Spodiopogon, Eulalia, Polytrias, Microstegium vimineum, Kerriochloa* and *Selima*. As this is proximal to the core Saccharinae (as compared with the Core Andropogoneae) by one argument this grouping should also be included within the Saccharinae subtribe. All other genera can be excluded as being either sister to the Core Andropogoneae or distal to them.

The ‘Saccharum complex’, as a potentially interbreeding group of species must be restricted to *Miscanthus, Miscanthidium, Pseudosorghum, Narenga, Erianthus s.s., Narenga*, trans-Himalayan species and *Saccharum*. In effect, the Saccharum complex hypothesis has been overturned and it has no validity, as least in terms of these species being involved in the direct evolution of *Saccharum*. However, the core species within the re-defined ‘Saccharum complex’ (which now correspond to the core species of the Saccharinae subtribe) may still be of interest to sugarcane breeders. It should also be noted that whilst hybridization and reticulation (network evolution) is common in the Andropogoneae as a whole, we find no evedence for reticulate evolution in the genera *Saccharum, Miscanthus* and *Miscanthidium*, though it has ocurred in *Narenga* and in the *Erianthus* species from the Americas and the trans-Himalayas.

## Materials and Methods

### Natural Language Processing

A proprietary natural language processing algorithm (Lloyd Evans and Joshi 2020) was employed to search for, index and mine text corpora (abstracts, full length papers, pre-prints, books, PhD thesis and on-line materials) for keyword combinations of genera and species plus the keywords phylogenetics, phylogenomics, phylogeny, taxonomy, relationships in all combinations. The subset of identified publications were read manually and meaningful data were extracted. Phylogenies identified in the publications (if not available on-line or in a database) were manually converted to Newick format.

### ITS-based Phylogeny

GenBank was searched with keywords to identify ITS sequences ¿ 500bp corresponding to ‘Core’ Andropogoneae, *Sorghum, Sarga, Saccharum, Miscanthus* and those species identified as lying between *Miscanthus* and *Saccharum. Tripidium* (*Erianthus* sect *Ripidium*) was employed as an outgroup. Longer sequences of 900bp (Snyman et al. 2018; Lloyd Evans and Joshi 2020) were employed as a backbone. The alignment was optimized as described previously (Martin et al. 2017) and a Maximum-Likelihood phylogeny was generated with IQ-Tree (Nguyen et al. 2015). Branch supports were derived as SH-aLRT single branch tests and non-parametric bootstrap with IQ-Tree as well as Bayesian Inference with Mr Bayes (Huelsenbeck, and Ronquist 2001). Phylogenetic trees were drawn with FigTree (http://tree.bio.ed.ac.uk/software/figtree/) and finished with Inkscape (https://inkscape.org/). Species names and voucher accessions along with NCBI accessions for the sequences employed in the phylogeny are given in Supplementary Table 1.

### Community Based Phylogeny

The phylogeny of Lloyd Evans et al. (2019) was employed as the backbone for the community tree. Additional phylogenies were integrated with the phangorn R framework (Schliep et al. 2016). Where possible, branch lengths of the backbone tree were retained and branch lengths of appended subtrees were scaled based on conserved common nodes. Individual nodes derived from academic literature were appended to the finished phylogeny using Archaeopteryx (Han & Zmasek 2009).

## References

Arthan W, McKain MR, Traiperm P, Welker CA, Teisher JK, Kellogg EA. 2017. Phylogenomics of Andropogoneae (Panicoideae: Poaceae) of Mainland Southeast Asia. Systematic Botany, 42:418–431.

Besse P, Taylor G, Carroll B, Berding N, Burner D, McIntyre CL. 1998. Assessing genetic diversity in a sugarcane germplasm collection using an automated AFLP analysis. Genetica, 104:143–153.

Clayton WD, Renvoize SA. 1986. Genera graminum. Grasses of the world. Genera graminum. Grasses of the World. 13.

Daniels J, Smith P, Paton N, Williams C.A. 1975. The origin of the genus Saccharum. Sugarcane Breeding News. 36:24–39.

D’Hont A, Souza GM, Menossi M, Vincentz M, van-Sluys MA, Glaszmann JC, Ulian E. 2008. Sugarcane: a major source of sweetness, alcohol, and bioenergy. In: Moore P.H, Ming R. editors. Plant genetics and genomics: crops and models. pp. 483–513 New York: Springer.

Dumortier BCJ. 1824. Observations sur les Graminées de la Flore Belgique. i-viii, [9]–153, t. 1–15

Estep MC, McKain MR, Diaz DV, Zhong J, Hodge JG, Hodkinson TR, Layton DJ, Malcomber ST, Pasquet R, Kellogg EA. 2014. Allopolyploidy, diversification, and the Miocene grassland expansion. Proceedings of the National Academy of Sciences, 111:15149–15154.

Grisebach AHR. 1846. Spicilegium florae rumelicae et bithynicae. 2:472

Grivet L, Daniels C, Glaszmann JC, D’Hont A. 2004. A review of recent molecular genetics evidence for sugarcane evolution and domestication. Ethnobotany Research and Applications, 2:009–017.

Guala GF. 1998. Revisions of Agenium and Homozeugos: integrating cladistic analysis and geographic information systems. Doctoral Dissertation. University of Florida: Gainesville.

Guala GF. 2000. The Relation of Space and Geography to Cladogenic Events in Agenium and Homozeugos (Poaceae: Andropogoneae) in South America and Africa. In Grasses: Systematics and Evolution: Systematics and Evolution, (SWL Jacobs, J Everett eds). CSIRO PUBLISHING.

Han MV, Zmasek CM. 2009. phyloXML: XML for evolutionary biology and comparative genomics BMC Bioinformatics, 10:356.

Hodkinson TR, Chase MW, Lledó DM, Salamin N, Renvoize SA. 2002. Phylogenetics of Miscanthus, Saccharum and related genera (Saccharinae, Andropogoneae, Poaceae) based on DNA sequences from ITS nuclear ribosomal DNA and plastid trnL intron and trnL-F intergenic spacers. Journal of plant research, 115:381–392.

Hoshino T, Davidse G. 1988. Chromosome numbers of grasses (Poaceae) from southern Africa. I. Ann. Missouri Bot. Gard. 75: 866–873.

Huelsenbeck JP, Ronquist F. 2001. MRBAYES: Bayesian inference of phylogenetic trees. Bioinformatics, 17:754–755.

Ibaragi Y, Kobayashi S. 2008. Veldkampia (Gramineae), a New Genus from Myanmar. Journal of Japanese Botany, 83:106.

Jensen KB, Highnight K, Wipff KJ. 1989. IOPB chromosome data 1. Int. Organ. Pl. Biosyst. Newslett. (Zurich) 13:20–21.

Kellogg EA, Birchler JA. 1993. Linking phylogeny and genetics: Zea mays as a tool for phylogenetic studies. Systematic Biology, 42:415–439.

Kellogg, E.A. and Watson, L., 1993. Phylogenetic studies of a large data set. I. Bambusoideae, Andropogonodae, and Pooideae (Gramineae). The Botanical Review, 59(4), pp.273–343.

Kellogg EA. 2013. Phylogenetic relationships of Saccharinae and Sorghinae. In Genomics of the Saccharinae. Springer, New York, NY. Pp 3–21.

Knuth KS. 1815. Considérations générales sur les Graminées. Mémoires du Muséum d’Histoire Naturelle 2:62–75.

Lloyd Evans D, Joshi SV. 2016. Complete chloroplast genomes of Saccharum spontaneum, Saccharum officinarum and Miscanthus floridulus (Panicoideae: Andropogoneae) reveal the plastid view on sugarcane origins. Systematics and Biodiversity. 14:548–571.

Lloyd Evans D, Joshi SV, Wang J. 2019. Whole chloroplast genome and gene locus phylogenies reveal the taxonomic placement and relationship of Tripidium (Panicoideae: Andropogoneae) to sugarcane. BMC evolutionary biology, 19:33.

Lloyd Evans D, Hughes B. 2020. Complete Chloroplast Genomes of *Saccharum giganteum, Saccharum longisetosum, Cleistachne sorghoides, Sarga timorense, Narenga porphyrocoma* and *Tripsacum dactyloides*. Comparisons with ITS phylogeny and Placement within *Saccharum*. BioRxiv. 10.1101/2020.06.12.149476

Martin LA, Lloyd Evans D, Castlebury LA, Sifundza JT, Comstock JC, Rutherford RS, McFarlane SA. 2017. Macruropyxis fulva sp. nov., a new rust (Pucciniales) infecting sugarcane in southern Africa. Australasian Plant Pathology, 46:63–74.

Mehra PN, Sharna ML. 1975. Cytological studies in some central and eastern Himalayan grasses. I. The Andropogoneae. Cytologia 40:61–74.

Mukherjee SK. 1954. Revision of the genus Saccharum. Bull. Bot. Soc. Bengal 8:143–148

Nair NV, Selvi A, Sreenivasan TV, Pushpalatha KN, Mary S. 2005. Molecular diversity amongSaccharum, Erianthus, Sorghum, Zea and their hybrids. Sugar Tech, 7:55–59.

Nguyen LT, Schmidt HA, Von Haeseler A, Minh BQ. 2015. IQ-TREE: a fast and effective stochastic algorithm for estimating maximum-likelihood phylogenies. Molecular biology and evolution, 32:268–274.

Piot A, Hackel J, Christin PA, Besnard G. 2018. One-third of the plastid genes evolved under positive selection in PACMAD grasses. Planta, 247:255–266.

Schliep K, Potts AA, Morrison DA, Grimm GW. 2016. Intertwining phylogenetic trees and networks (No. e2054v1). PeerJ Preprints.

Selvi A, Nair NV, Noyer JL, Singh NK, Balasundaram N, Bansal KC, Koundal KR, Mohapatra T. 2006. AFLP analysis of the phenetic organization and genetic diversity in the sugarcane complex, Saccharum and Erianthus. Genetic Resources and Crop Evolution, 53:831–842.

Setshogo MP. 1997. TAXONOMIC STUDIES AND GENERIC DELIMITATION IN THE GRASS SUBTRIBE Sorghinae. PhD Thesis, University of Edinburgh.

Skendzic EM, Columbus JT, Cerros-Tlatilpa R. 2007. Phylogenetics of Andropogoneae (Poaceae: Panicoideae) based on nuclear ribosomal internal transcribed spacer and chloroplast trnL–F sequences. Aliso: A Journal of Systematic and Evolutionary Botany, 23:530–544.

Snyman SJ, Komape D, Khanyi H, Van Den Berg J, Cilliers D, Lloyd Evans D, Barnard S, Siebert SJ (2018) Assessing the likelihood of gene flow from sugarcane (Saccharum hybrids) to wild relatives in South Africa. Frontiers in Bioengineering and Biotechnology, 6:72.

Soreng RJ, Peterson PM, Romaschenko K, Davidse G, Teisher JK, Clark LG, Barberá P, Gillespie LJ, Zuloaga FO. 2017. A worldwide phylogenetic classification of the Poaceae (Gramineae) II: An update and a comparison of two 2015 classifications. Journal of Systematics and Evolution, 55:259–290.

Sreenivasan TV, Sreenivasan J. 1984. Cytology of Saccharum complex from New Guinea, Indonesia and India. Caryologia 37:351–357.

Tiwari AP, Chorghe A. 2019. Lectotypification of the monotypic genus Pseudodichanthium (Andropogoneae: Poaceae). Phytotaxa, 415:150–152.

Welker CA, Souza-Chies TT, Longhi-Wagner HM, Peichoto MC, McKain MR, Kellogg EA. 2015. Phylogenetic analysis of Saccharum sl (Poaceae; Andropogoneae), with emphasis on the circumscription of the South American species. American Journal of Botany, 102:248–263.

Zachos FE. 2016. Species concepts in biology. Historical Development, Theoretical Foundations and Practical Relevance. Springer. NY.

