## Supplementary Document 1 for "On the Validity of the Saccharum Complex and the Saccharinae Subtribe: A Re-assesment"

| Species | Voucher Accession | GenBank Accession |
|---|---|---|
| *Sorghum nitidum* | O. Neamsuvan 216 | GQ856354.1 |
| *Sorghum timorense* | | GQ121728.1 |
| *Sorghum versicolor* | WSVE1 | KT933739.1 |
| *Saccharum spontaneum* | IS-76-164 | AB281145.1 |
| *Saccharum spontaneum* | SES-597 | AB281147.1 |
| *Saccharum robustum* | R13 | AF345239.1 |
| *Saccharum officinarum* | Mangeer | AB250692.1 |
| *Saccharum hybrid* | CP58-48.1 | KX495638.1 |
| *Miscanthus junceus* | genotype 89 | AY116289.1 |
| *Miscanthus junceus* | genotype 88 | AY116288.1 |
| *Miscanthus ecklonii* | Genotype 86 | AY116290.1 |
| *Saccharum baldwinii* | Columbus 4181 | DQ005049.1 |
| *Saccharum brevibarbe var. contortum* | genotype 121 | AY116287.1 |
| *Miscanthus fuscus* | genotype 83 | AY116286.1 |
| *Narenga porphyrocoma* | R33 | AF345233.1 |
| *Narenga porphyrocoma* | Zeng 0381 | JX156343.1 |
| *Erianthus rockii* | isolate R38 | AF345217.1 |
| *Erianthus rockii* | isolate R37 | AF345216.1 |
| *Miscanthus nudipes* | Dn-ITS-94f | KM241751.1 |
| *Miscanthus nepalensis* | HN3176 | JN544347.1 |
| *Miscanthus nepalensis* | genotype 25 | AY116292.1 |
| *Miscanthus nudipes* | HN3178 | JN544346.1 |
| *Narenga fallax* | | AF345213.1 |
| *Erianthus fulvus* | R40 | AF345218.1 |
| *Erianthus fulvus* | R42 | AF345220.1 |
| *Pseudosorghum zollingeri* | J.D. Wogch 35030 (L) | GQ856357.1 |
| *Pseudosorghum fasciculare* | B. Balansa 1770 (L) | GQ856355.1 |
| *Miscanthus paniculatus* | HN3181 | JN544345.1 |
| *Miscanthus sacchariflorus var. gracilis* | KWNU75843 | HQ822019.1 |
| *Miscanthus lutarioriparius* | | MK138898.1 |
| *Miscanthus purpurascens* | HN2437 clone 1 | JN544332.1 |
| *Miscanthus purpurascens* | HN2393 clone 2 | JN544330.1 |
| *Miscanthus lutarioriparius* | A0105 | MK981299.1 |
| *Miscanthus sacchariflorus* | sn2727 | AJ426564.1 |
| *Miscanthus x giganteus* | S1a | KU297965.1 |
| *Miscanthus sacchariflorus* | KWNU75842 | HQ822018.1 |
| *Miscanthus changii* | EWH9013 | HQ822028.1 |
| *Miscanthus tinctorius* | TI_1 | LC113999.1 |
| *Miscanthus tinctorius* | TI_2 | LC114000.1 |
| *Miscanthus intermedius* | IN_1 | LC113995.1 |
| *Miscanthus x giganteus* | 1780, isolate 2 | AJ426563.1 |
| *Miscanthus oligostachyus* | KWNU75835 | HQ822027.1 |
| *Miscanthus floridulus* | | MK138896.1 |
| *Miscanthus sinensis* | Q561 | MH711442.1 |
| *Miscanthus transmorrisonensis* | genotype 65 | AY116271.1 |
| *Miscanthus transmorrisonensis* | MstT-ITS-87f | KM241678.1 |
| *Miscanthus floridulus* | C0128 | MK981287.1 |
| *Miscanthus purpurascens* | HN2437 | JN544333.1 |

| | | |
|---|---|---|
| *Miscanthus x giganteus* | 1780, isolate 1 | AJ426562.1 |
| *Miscanthus sinensis* | HN2457 | JN544315.1 |
| *Miscanthus purpurascens* | HN2438 | JN544335.1 |
| *Miscanthus purpurascens* | HN2393 clone 1 | JN544331.1 |
| *Miscanthus sinensis* | HN2401 | JN544321.1 |
| *Miscanthus floridulus* | Zeng 0415 | JX156342.1 |
| *Miscanthus oligostachyus* | genotype 161 | AY116279.1 |
| *Sorghastrum nutans* | Skendzic 5065 RSA | DQ005080.1 |
| *Sorghastrum secundum* | Columbus 4243 RSA | DQ005078.1 |
| *Ischaemum afrum* | | HM347038.1 |
| *Spodiopogon cotulifer* | HCCN-PJ008548-PB-339 | KF163703.1 |
| *Spodiopogon sibiricus* | Q560 | MH711441.1 |
| *Themeda quadrivalvis* | MSV1038 | KY991095.1 |
| *Themeda triandra* | HCCN-PJ008548-PB-214 | KF163704.1 |
| *Heteropogon contortus* | LSC101 | MH768196.1 |
| *Heteropogon contortus* | TuTY1505 | MH768197.1 |
| *Heteropogon triticeus* | EPA3355 | KY991073.1 |
| *Andropogon glomeratus var. scabriglumus* | CBP069 | MF964041.1 |
| *Diheteropogon amplectens* | | KY991078.1 |
| *Cymbopogon flexuosus* | CF-9 | KX828251.1 |
| *Cymbopogon flexuosus* | CF-10 | KX828252.1 |
| *Schizachyrium brevifolium* | HCCN-PJ008548-PB-256 | KF163598.1 |
| *Schizachyrium condensatum* | C. Silva 843 | KP878911.1 |
| *Eulalia villosa* | genotype 132 | AY116302.1 |
| *Bothriochloa ischaemum* | TuTY1522 | MH768166.1 |
| *Microstegium vimineum* | HCCN-PJ008548-PB-284 | KF163837.1 |
| *Microstegium vimineum* | HCCN-PJ008548-PB-344 | KF163649.1 |
| *Sehima nervosum* | | GQ870205.1 |
| *Kerriochloa siamensis* | | GQ870204.1 |
| *Ischaemum anthephoroides* | HCCN-PJ008548-PB-253 | KF163647.1 |
| *Sorghum laxiflorum* | | GQ121741.1 |
| *Sorghum macrospermum* | | GQ121742.1 |
| *Sorghum nitidum* | Zeng 0367 | JX156346.1 |
| *Sorghum bicolor* | FS035 | MH762138.1 |
| *Saccharum perrieri* | TAN:NOP140 | MN342165.1 |
| *Lasiorhachis hildebrandtii* | TAN:LRK2008 | MN342164.1 |
| *Saccharum arundinaceum* | IND-84-394 | AB281160.1 |
| *Saccharum arundinaceum* | Zeng 0407 | JX156345.1 |
| *Erianthus ravennae* | | AF019824.1 |
